# Pharmacodynamic Markers of LRRK2 Inhibition in Biofluids

**DOI:** 10.1101/2020.01.28.923557

**Authors:** Shijie Wang, Kaela Kelly, Nathalie Schussler, Sylviane Boularand, Laurent Dubois, Frank Hsieh, Elizabeth Tengstrand, Jonathan M. Brotchie, James B. Koprich, Andrew B. West

## Abstract

Hyper-activated LRRK2 is linked to Parkinson’s disease susceptibility and progression. Quantitative measures of LRRK2 inhibition, especially in the brain, may be critical in the clinical development of successful LRRK2-targeting therapeutics. In this study, three different brain-penetrant and selective LRRK2 small-molecule kinase inhibitors (PFE-360, MLi2, and RA283) were orally administered to groups of cynomolgus macaques at different doses. Biofluid markers with proposed pharmacodynamic properties for assessing LRRK2 inhibition were measured from samples of blood, urine, and cerebral-spinal fluid (CSF). LRRK2 kinase inhibition led to consistent reduced pS935-LRRK2 and pRab10 proteins in blood mononuclear cells, reduced exosome LRRK2 protein and di-docosahexaenoyl (22:6) bis (monoacylglycerol) phosphate in urine, and reduced exosome LRRK2 and autophosphorylated pS1292-LRRK2 protein in CSF. Incomplete LRRK2 kinase inhibition reduced LRRK2 protein secretion in exosomes whereas high drug exposures may reduce both exosome and tissue levels of LRRK2 protein. These orthogonal markers for LRRK2 inhibition in urine and CSF can be used in combination with blood markers to non-invasively monitor the potency of LRRK2-targeting therapeutics in the brain and periphery.

## Introduction

Leucine-rich repeat kinase 2 (LRRK2) small molecule kinase inhibitors and anti-sense oligonucleotides have demonstrated neuroprotective efficacy in some models of Parkinson’s disease (PD) and represent a promising novel class of disease modifying therapeutics for neurodegeneration (Daher et al., 2015;Volpicelli-Daley et al., 2016;Zhao et al., 2017). Efforts to develop potent and selective therapies that mitigate LRRK2-hyperactivation associated with pathogenic *LRRK2* mutations have recently intensified in industry (West, 2017;Alessi and Sammler, 2018). Reliable measures of LRRK2 expression and enzymatic inhibition may assist in the identification of the most efficacious LRRK2-targeting therapies at optimized doses, particularly in early clinical trials. As some LRRK2-targeting molecules have recently progressed through initial safety trials (Therapeutics, 2018), non-invasive pharmacodynamic measures in biofluids may be valuable to establish on-target drug effects in the brain and other tissues. With a renewed emphasis on precision approaches in the clinic, comprehensive demonstrations of robust target engagement during initial safety trials (e.g., in acute ascending dose strategies) may help successfully identify the best molecules to transition to larger efficacy studies in limited populations of genetically-defined LRRK2-linked PD (Ivy et al., 2010;Sistare et al., 2010;West, 2017).

LRRK2 is a phospho-protein in all cells and tissues so-far evaluated (West et al., 2007), with a cluster of phosphorylated residues controlled by several other kinases and phosphatases that can regulate LRRK2 interactions with 14-3-3 proteins (Dzamko et al., 2010;Nichols et al., 2010;Li et al., 2011). Small molecule LRRK2 kinase inhibitors acutely reduce constitutive phospho-sites in LRRK2, most notably pS935 (Dzamko et al., 2010;Nichols et al., 2010), as well as the autophosphorylated residue pS1292 located in the C-terminal region of the LRRK2 leucine-rich repeat (LRR) domain, nearby the Rab-like ROC domain (Henry et al., 2015;Wang et al., 2017;Liu et al., 2018). LRRK2 robustly autophosphorylates its own Rab-like GTPase domain *in vitro* (Greggio et al., 2009;Muda et al., 2014;Liu et al., 2016), in addition to phosphorylating other GTP-bound Rab proteins (i.e., Rab10) both *in vitro* and *in vivo* (Steger et al., 2016;West and Cookson, 2016). LRRK2 kinase inhibitors reduce levels of phospho-Rab10 in *ex vivo* inhibitor treatments of immune cells that express high levels of LRRK2 protein (e.g., monocytes, neutrophils, and whole-lysates of mononuclear cells), as well as in tissues (e.g., lung and kidney) procured from mice and rats (Fan et al., 2018;Kelly et al., 2018;Lis et al., 2018). pS1292-LRRK2 and pT73-Rab10 are direct substrates of LRRK2, although the phosphatases that affect these sites are likely different in cells and tissues, potentially affecting pharmacodynamic marker kinetics (Berndsen et al., 2019). Apart from protein pharmacodynamic markers, levels of di-docosahexaenoyl (22:6) bis (monoacylglycerol) phosphate (di-22:6-BMP) which are usually elevated in phospholipidosis and lysosomal storage diseases (Meikle et al., 2008;Liu et al., 2014;Tengstrand et al., 2019), appear decreased in urine from LRRK2-inhibitor treated non-human primates (Fuji et al., 2015). The mechanisms behind LRRK2-inhibitor induced decreases in di-22:6-BMP in urine have yet to be defined.

Recently we identified LRRK2 protein secreted by cells in a type of microvesicle called exosomes that are derived from multivesicular bodies (Fraser et al., 2013). Although only a small fraction of LRRK2 protein is secreted into exosomes, the ratio of phospho-LRRK2 to total LRRK2 levels appears comparable between exosome and total cell (or tissue) lysates (Wang et al., 2017). The exosome fraction of biofluids (sometimes called liquid-biopsies) allows the assessment of cellular levels of proteins from tissues that are otherwise inaccessible for analysis, for example the brain. LRRK2 protein is widely expressed through the body, with inducible expression in some immune cell subsets with pro-inflammatory agonists, and constitutive tissue expression in some cells in the brain, lung, and kidney (Biskup et al., 2007;Maekawa et al., 2010;Hakimi et al., 2011;Fuji et al., 2015;West, 2017). In some model systems, LRRK2 kinase inhibitors, usually in great excess of binding affinity concentrations, can reduce total LRRK2 protein levels comparably to effects seen with anti-sense LRRK2-targeting oligonucleotides (Skibinski et al., 2014;Zhao et al., 2017). The reduction of total LRRK2 protein in cells and tissues may relate to how specific inhibitors interact in the ATP-pocket, whereas LRRK2 inhibitor-induced reductions in exosome secretion in biofluids may relate to LRRK2 subcellular localization changes in the endolysosomal compartment broadly caused by kinase inhibition (Fraser et al., 2013;Lobbestael et al., 2016;Kelly et al., 2018;De Wit et al., 2019). Although phospho-LRRK2 to total-LRRK2 ratios have been suggested as efficacious measures of LRRK2 inhibitors (Dzamko et al., 2010), inhibitor-induced destabilization of LRRK2 protein or reductions of secreted LRRK2 protein may confound interpretation of these ratios.

Biochemical surveys of LRRK2 kinase activity in *ex vivo* assays of proteins purified from mouse-derived cells and tissues reveal much higher LRRK2 activity in the brain than in other tissues and cells (Li et al., 2010). Further, autophosphorylated LRRK2 protein (i.e., pS1292-LRRK2) appears particularly high in human CSF exosomes as compared to exosomes procured from human urine or lysates from immune cells (Wang et al., 2017). Non-invasive pharmacodynamic markers to measure LRRK2 inhibition in the brain are sought after but have not been previously described. Herein we better define the relationship between LRRK2 inhibition and different proposed pharmacodynamic markers, focusing on the clinically available biofluids in blood, urine, and CSF. We employ distinct, orally available, highly potent (low-to-sub nanomolar-binding), highly-selective, small molecule LRRK2 kinase inhibitors PFE-360, MLi2, and RA283 to ascertain variability associated with each pharmacodynamic marker at both baseline and with drug treatment (Fell et al., 2015;Kelly et al., 2018). The use of non-human primates that have pharmacokinetic responses more similar to humans compared to rodents further allows the collection of sufficient volumes of urine and CSF for LRRK2 detection at the same time as blood collections to identify markers with the potential for translation to human studies.

## Materials and Methods

### Compounds

Small-molecule LRRK2 kinase inhibitors PFE-360, MLi2, and RA283, were synthesized in-house. The identity and high purity (i.e., >97.4% for each compound) of each compound was confirmed by nuclear-magnetic resonance and mass spectrometry. PFE-360 was originally discovered as part of a Pfizer chemistry program (Henderson et al., 2015), MLi2 was originally discovered as part of a Merck chemistry program (Fell et al., 2015;Scott et al., 2017), and RA283 was discovered as part of a Sanofi chemistry program.

Previously assessed *in vitro* kinase assays show similar IC_50_ potencies with the three structurally distinct inhibitors, with PFE-360 demonstrating 6 nM IC_50_ inhibition for WT-LRRK2 autophosphorylation and MLi2 demonstrating 4 nM inhibition in transfected HEK293T assessments (Kelly et al., 2018). RA283 demonstrates an IC_50_ of 15 nM on WT-LRRK2 activity in transfected HEK293T cells.

Selectivity profiling of over 300 protein kinases with each of these molecules in different assay platforms reveals high selectivity. Off-target inhibition (kinases other than LRRK2) with IC_50_< 500 nM identifies two other kinases (MST-2, MAP3K5) for PFE-360 (Thirstrup et al., 2017) and three other kinases for MLi2 (MAP3K14, CLK4, CLK2)(Fell et al., 2015;Scott et al., 2017). The kinase selectivity of RA283 was assessed using the Eurofin panel where two kinases besides LRRK2 of >300 tested were inhibited with IC_50_ <500 nM (MLK1 108 nM; ACK1 188 nM).

### Non-human primate (NHP) treatment

For the PFE-360 and MLi2-treated groups, female cynomolgus macaques, with an average age of 10 years (Xishan, PRC) were orally administered 5 mg kg-1 PFE-360 or MLi2 daily for five days (N = 5 per group, 10 total). Under anesthesia (Zoletil/atropine (6/0.04 mg/kg, IM), blood, urine (via acute catheterization), and CSF (from cisterna magna) were collected before treatment (baseline) and two hours post final treatment for a longitudinal assessment. Indices of LRRK2 inhibition were comparisons of baseline levels to those levels after drug-treatment.

For the RA283 treated groups, male cynomolgus macaques, ∼3.5 years (Tamarinier and Noveprim), were orally administered RA283 at a low dose (10 mg kg^-1^) and a high dose (30 mg kg^-1^) regimen for seven days together with a vehicle-only group (n=2 per group, 6 total). All procedures were performed in full compliance with standards for the care and use of laboratory animals, according to French and European Community (Directive 2010/63/EU) legislation, and approved by the local Animal Ethics Committee and the French Ministry for Research, under authorization H94-002-4 arrêté 2015-160 (10 December 2015). Urine, blood and CSF samples were collected 2 and 6 hours post final treatment for the low dose and high dose RA283 groups respectively. Indices of LRRK2 inhibition were comparisons of levels in the vehicle-only group to those in the low- and high-dose groups.

### Drug plasma measurements

Unbound drug plasma concentrations were quantified by LC/MS analysis as previously described (Kelly et al., 2018).

### Exosome isolation

Urine (∼200 µL to 1 mL) and CSF samples (∼200 µL to 1 mL) were stored at -80°C in cryo-vials (Corning) and quick thawed in a water bath at 42°C before processing. Fluids were centrifuged at 10,000 x g for 30 minutes and the supernatants were transferred to polycarbonate centrifuge tubes. Samples were then centrifuged at 150,000 x g for 2 hours at 4°C. Supernatants were used for di-22:6-BMP analyses (below). Pellets (enriched in exosomes) were lysed in Laemmli buffer (4% SDS, 10% glycerol, 120 mM Tris pH 6.8, 40 mM NaF) freshly supplemented with 50 mg mL-1 DTT (dithiothreitol), and stored at -80°C. Consistent with past approaches, TSG101 was selected as an exosome housekeeping protein and loading control in urinary exosomes, whereas flotillin-1 was selected as an exosome housekeeping protein and loading control in CSF-derived exosomes, owing to the relative differential abundance of each protein in the respective biofluid (Wang et al., 2017).

### Protein measurements

Peripheral blood mononuclear cells (PBMCs) were isolated from whole blood and lysed in Laemmli buffer (described above). Frontal cortices (300 mg) were lysed in Precellys tubes (CK14) in 1 ml RIPA buffer (1x, Cell Signaling) supplemented with phosphatase inhibitors (okadaic acid 1 µM and sodium fluoride 100 mM) and protease inhibitors (Complete cocktail, Sigma). After one-hour incubation at 4°C, lysates were centrifuged for 10 min at 4°C at 15,000 x g. Primary antibodies (all used at 1:1000, or ∼0.5 to 1.0 μg mL^-1^) used include anti-LRRK2 (N241A/34, Antibodies Inc), anti-LRRK2 (clone MJFF2 c41-2, Abcam), anti-pS935 LRRK2 (clone UDD2-10, Abcam), anti-pS1292-LRRK2 (clone MJFR-19-7-8, Abcam), anti-pT73 Rab10 antibody (clone MJF-R21, Abcam), anti-Rab10 (clone D36C4, Cell Signaling), anti-flotillin-1 (clone D2V7J, Cell Signaling), and anti-TSG101 (ab30871, Abcam). The following secondary antibodies (1:10,000) were used: HRP conjugated donkey anti-rabbit secondary antibody (Jackson ImmunoResearch, # 711-035-152) and HRP conjugated donkey anti-mouse secondary antibody (Jackson ImmunoResearch, # 715-035-151). Immunoblots were developed using Luminata Crescendo Western HRP Substrate and digital signals were imaged and analyzed using ChemiDoc Imaging system (Bio-Rad) and ImageLab software (BioRad). Additionally, a homogeneous time-resolved fluorescence (HTRF) immunoassay was used for LRRK2 and pS935-LRRK2 according to manufacturer’s guidelines (Cisbio). Recombinant full-length human G2019S-LRRK2 protein (Invitrogen) was used as a protein standard. Assay linearity and variably associated with each measure in the different biofluids was previously reported (Fraser et al., 2016;Wang et al., 2017;Kelly et al., 2018). Standard curves with recombinant protein demonstrated linearity between 10 pg to 500 pg for total LRRK2 and 2.8pg to 140pg for pS1292-LRRK2 to (r>0.9) as shown previously (Wang et al., 2017). Recombinant phospho-Rab proteins were not available for this study. Repeated measure analysis of four biofluid samples from NHPs limited to ∼200 µL of starting biofluid volume showed a coefficient of variation (CV) better than 20%.

### Di-docosahexaenoyl (22:6) bis (monoacylglycerol) phosphate measurement

Di-docosahexaenoyl (22:6) bis (monoacylglycerol) phosphate (di-22:6-BMP) was measured in plasma and exosome-depleted urine and CSF by ultra-performance liquid chromatography – tandem mass spectrometry (UPLC-MS/MS). The analyses were performed as previously described (Tengstrand et al., 2019). Standard curves were prepared using an authentic di-22:6-BMP reference standard. Di-22:6-BMP-d5 was used as an internal standard. Analyses were conducted using a Nextcea XR UPLC system (Shimadzu). A SCIEX X500B QTOF mass spectrometer with SCIEX OS software was used for quantitation (SCIEX). The calibration curve for di-22:6-BMP was linear between 0.05-100 ng mL-1 with ≥75% of the standards within ±15% (±20% for the lower-limit of quantitation) of the spiked concentration. Results at four quality-control concentrations (lower-limit, low, median, and high) showed that the assay was reproducible within ±15% of the nominal value (±20.0% at the lower-limit of quantitation). The concentrations of di-22:6-BMP measured in exosome-depleted urine were normalized to urine creatinine and reported as ng per mg of creatinine. Di-22:6-BMP values in exosome-depleted CSF were normalized by total protein and reported as ng per mg total protein.

### Statistical Analysis

Two-tailed, paired t-tests were used to compare baseline and LRRK2 inhibitor treated samples. The median baseline was calculated for each marker and set to 1.0. Statistical analyses were performed using GraphPad Prism 5.0. When total protein was significantly impacted by treatment (i.e., treatment-induced reduction of total LRRK2 protein), the ratio of phospho to total-protein was considered critically confounded and interpreted with caution.

## Results

To measure pharmacodynamic responses related to LRRK2 kinase inhibition in macaques, PBMCs, plasma, urine, and CSF were procured from ten animals prior to drug administration. Two groups of five macaques were then treated daily with either 5 mg kg^-1^ of PFE-360 or MLi2 (Supplemental Figure 1A, B). On the fifth day of treatment, PBMCs, plasma, urine, and CSF were collected two hours after the final drug dose to allow for a range of nanomolar plasma concentrations of unbound drug. Concentrations of PFE-360 varied in the five macaques at the time of collection from ∼40 nM to ∼145 nM. Plasma concentrations of MLi2 varied between ∼12 nM and ∼70 nM. No adverse drug reactions were noted in any animal.

**Figure 1.**
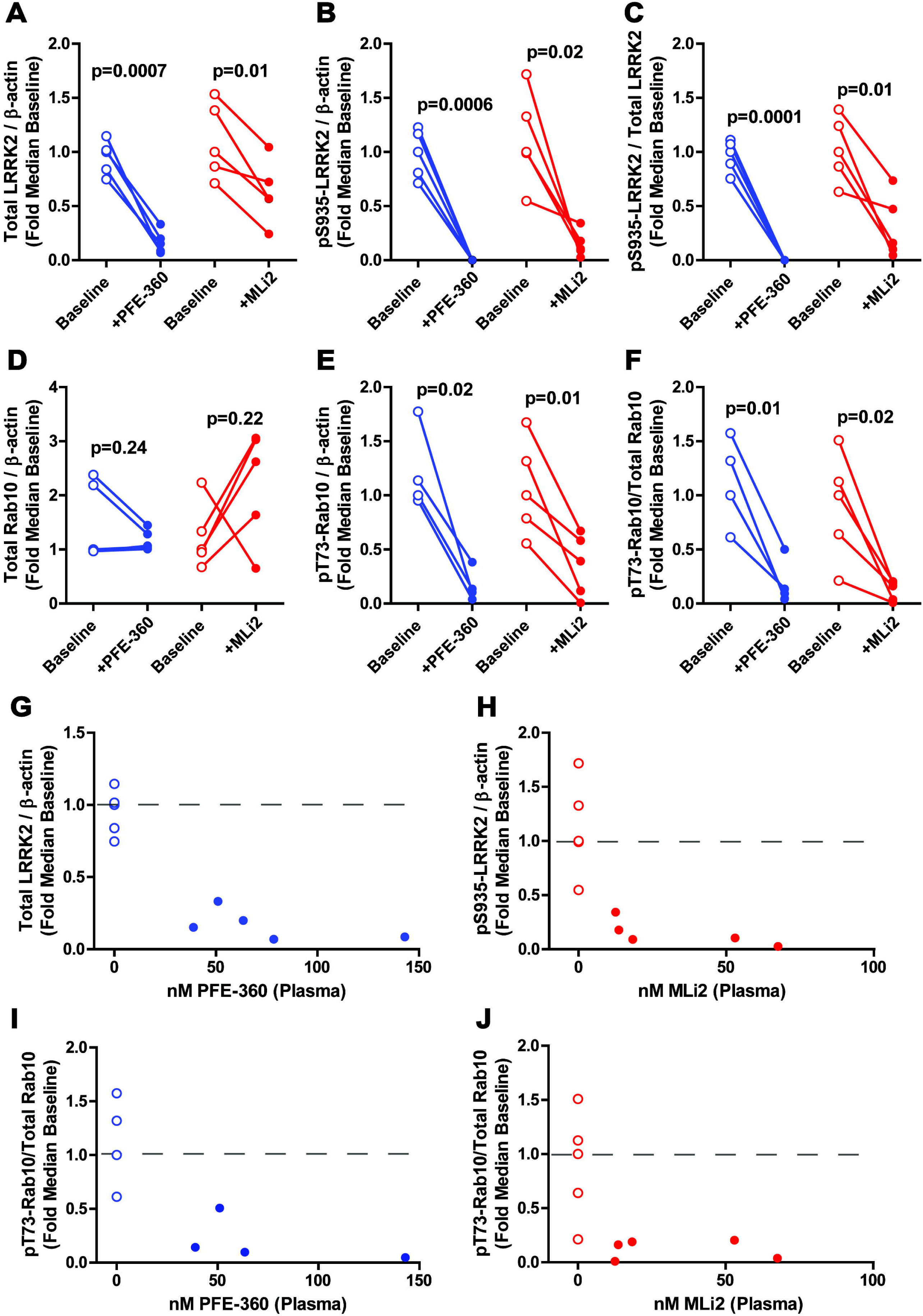
LRRK2 inhibition markers in PBMCs. Inhibition values for measures of LRRK2 and Rab10 in isolated PBMCs after five days of PFE-360 (blue) and MLi2 (red) treatment (5 mg kg^-1^ BID). Protein levels were normalized to the housekeeping protein β-actin. Fold median baseline values before and after (2-hours) treatment for **A**. Total LRRK2, **B**. pS935-LRRK2, **C**. pS935-LRRK2/Total LRRK2, **D**. Total Rab10, **E**. pT73-Rab10, and **F**. pT73-Rab10**/**Total Rab10. For Rab10 inhibition measures (panel D-F), one outlier sample in the PFE-360 treatment group was not included because baseline pT73-Rab10 levels fell below the limit of detection. Paired t-tests were performed to determine significance between baseline and post-drug exposures. Scatterplots of **G**. PFE-360 (nM) and total LRRK2, **H**. MLi2 (nM) and pS935-LRRK2, **I**. PFE-360 (nM) and pT73-Rab10/Total Rab10, and **J**. MLi2 (nM) and pT73-Rab10/Rab10. The dashed grey lines represent the median value with no drug (i.e., 0 nM). Open circles were used to represent baseline levels of each NHP. *p*-values were calculated using a paired t-test.

### Blood markers of LRRK2 kinase inhibition

LRRK2 is differentially expressed in immune cells in PBMCs, with some of the highest LRRK2 expression in monocytes, B-cells, and neutrophils (Kubo et al., 2010;Hakimi et al., 2011;Thévenet et al., 2011;Cook et al., 2017). Compared to the baseline levels of total LRRK2 in PBMCs, PFE-360 treatment caused a large drop in LRRK2 levels (normalized to β-actin, unadjusted for immune cell subtypes), with just 22% average of total LRRK2 protein remaining (Mdn_PFE-360_= 0.15, IQR= 0.07- 0.27, *t*(4)= 9.45, *p*=.0007), and undetectable quantities of pS935-LRRK2 (Figure 1 and Supplemental Figure 2A). The LRRK2 autophosphorylation site pS1292-LRRK2, detectable in human CSF and urinary exosomes in past studies (Wang et al., 2017;Wang et al., 2019), was not reliably detected in PBMC lysates. Total Rab10 levels, also normalized to β-actin and unadjusted for immune cell subtypes, were variable in expression in PBMCs (Mdn_PFE-360_= 1.07, IQR= 1.03- 1.37, *t*(4)= 1.37, *p*=.24, Figure 1D). Four of five macaques demonstrated a reduction in pT73-Rab10 levels (Mdn_PFE-360_= 0.12, IQR= 0.06- 0.32, Figure 1E, F). One outlying macaque had baseline levels of pT73-Rab10 that fell below our limit of detection (Supplemental Figure 2A, lane 1) and was excluded. Plasma levels of the phospholipid di-22:6-BMP were not reduced through PFE-360 treatments and varied in concentration, with an average of 8.07 ± 2.35 ng mL-1 at baseline and 7.50 ± 3.70 ng mL-1 after treatment.

**Figure 2.**
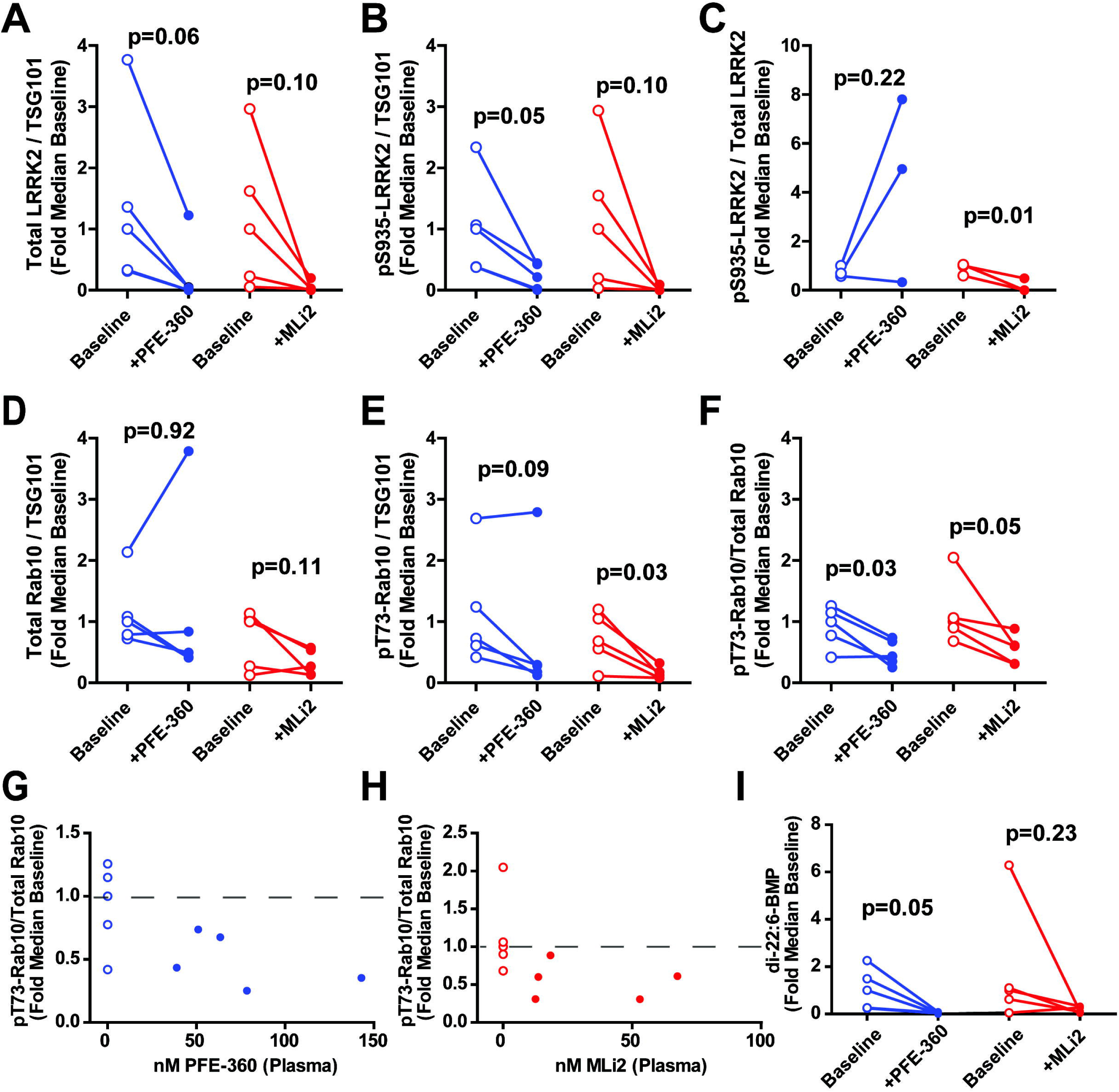
LRRK2 inhibition markers in urine. Inhibition values for measures of exosomal LRRK2 and Rab10 after five days of PFE-360 (blue) and MLi2 (red) treatment (5 mg kg^-1^ BID). Proteins isolated from urinary exosomes were normalized to the exosome marker TSG101. Fold median baseline values before and post-treatment for **A**. total LRRK2, **B**. pS935-LRRK2, and **C**. pS935-LRRK2/total LRRK2. Note, the ratio for two animals could not be plotted due to the total LRRK2 value recorded as zero. **D**. Total Rab10, **E**. pT73-Rab10, and **F**. pT73-Rab10**/**total Rab10, with paired t-tests to determine significance between baseline and post-drug exposure measures for PFE-360 and MLi2, respectively. Scatterplots between unbound plasma concentrations of **G**. PFE-360 (nM) or **H**. MLi2 (nM) and inhibition of pT73-Rab10/Rab10. The dashed grey lines represent the median value with no drug. **I**. Di-22:6-BMP levels (ng mg^-1^ Creatinine) in exosome-depleted urine at baseline and post-drug exposure of PFE-360 (blue) or MLi2 (red). Open circles were used to represent baseline levels of each NHP. *p*-values were calculated using a paired t-test.

In MLi2 treated macaques, total LRRK2 protein also diminished in PBMC pellets normalized to β-actin concentrations (Mdn_MLi2_= 0.57, IQR= 0.40- 0.88 *t*(4)=4.38, *p*= .01), but to a lesser extent than PFE-360, 47% diminished compared to 78% with PFE-360 on average (Figure 1A and Supplemental Figure 2B), consistent with our past experience with these drugs in rats and mice (Kelly et al., 2018). pS935-LRRK2 signal was detected in all MLi2-treated macaques at low levels compared to baseline (Mdn_MLi2_= 0.11, IQR= 0.06- 0.26, *t*(4)= 3.98, *p*=.02), although one macaque showed only a 37% reduction in pS935-LRRK2 that also had the lowest concentration of unbound plasma MLi2 (12.5 nM, Figure 1B and 1H). pT73-Rab10 levels reduced by 71% on average (Mdn_MLi2_= 0.39, IQR= 0.06- 0.63, *t*(4)= 4.35, *p*=.01, Figure 1E), even in the macaque with just 12.5 nM of drug in circulation. The ratio of pT73-Rab10 to total Rab10 protein offered a comparable indication to pT73-Rab10 measures alone (Mdn_MLi2_= 0.16, IQR= 0.02- 0.20, *t*(4)= 3.85, *p*=.02, Figure 1F, J). Overall, these results suggest that pS935-LRRK2 and pT73-Rab10 levels are reliably reduced when normalized to β-actin in PBMCs after LRRK2 kinase inhibitor treatment, whereas total LRRK2 levels and total Rab10 levels were more variably affected.

### Urine markers of LRRK2 kinase inhibition

Our previous work localized LRRK2 protein expression in urine to urinary exosomes (Fraser et al., 2016;Wang et al., 2019). Compared to baseline measures of LRRK2 in macaque urinary exosomes, PFE-360 or MLi2 treatment caused a profound decrease in total LRRK2 protein normalized to the exosome house-keeping protein TSG101 in all NHPs, although the variability at baseline measures was higher than the measures observed in PBMCs, with two NHPs in the each group presenting with very low LRRK2 levels at baseline that affected group comparisons (Mdn_PFE-360_= 0.04, IQR= 0- 0.64, *t*(4)= 2.66, *p*=.06; Mdn_MLi2_= 0.02, IQR= 0.01- 0.11, *t*(4)= 2.15, *p*=.10, Figure 2A and Supplemental Figure 3). Comparable variabilities were noted with pS935- LRRK2 protein (Mdn_PFE-360_= 0.21, IQR= 0.01- 0.43, *t*(4)= 2.84, *p*=.05; Mdn_MLi2_= 0, IQR= 0- 0.04, *t*(4)= 2.15, *p*=.10, Figure 2B and Supplemental Figure 3). Owing to the reductions in total LRRK2 protein caused by kinase inhibition, the ratio of pS935-LRRK2 to total LRRK2 protein was uninformative, especially in animals where total LRRK2 protein was most dramatically reduced (Figure 2C). Similar to PBMCs, the LRRK2 autophosphorylation site pS1292 was not reliably detected in macaque urine in our assay. Total Rab10 levels, also normalized to the exosome housekeeper TSG101, did not vary with drug treatment (Mdn_PFE-360_= 0.50, IQR= 0.41- 2.31, *t*(4)= 0.10, *p*=.92; Mdn_MLi2_= 0.27, IQR= 0.14- 0.56, *t*(4)= 2.03, *p*=.11, Figure 1D). Decreases were observed with pT73-Rab10 in most macaques (Mdn_PFE-360_= 0.28, IQR= 0.15- 1.54, *t*(4)= 2.25, *p*=.09; Mdn_MLi2_= 0.12, IQR= 0.08- 0.25, *t*(4)= 3.30, *p*=.03, Figure 2E). The ratio of pT73-Rab10 to total Rab10 protein decreased particularly in the macaques with the highest plasma concentrations of PFE-360 (∼145 nM, 50% remaining from baseline, Mdn_PFE-360_= 0.43, IQR= 0.30- 0.71, *t*(4)= 3.46, *p*=.03) and MLi2 (∼70 nM, 30% remaining from baseline, Mdn_MLi2_= 0.60, IQR= 0.31- 0.75, *t*(4)= 2.70, *p*=.05, Figure 2F, G, H). Overall, PFE-360 and MLi2 treatment reduced total LRRK2 protein secretion in urinary exosomes, although the reduction was more variable in macaques that had low baseline levels of LRRK2. As with PBMC analyses, ratios of phospho- to total protein as a pharmacodynamic measure could be considered confounded due to variable changes of total LRRK2 protein levels.

**Figure 3.**
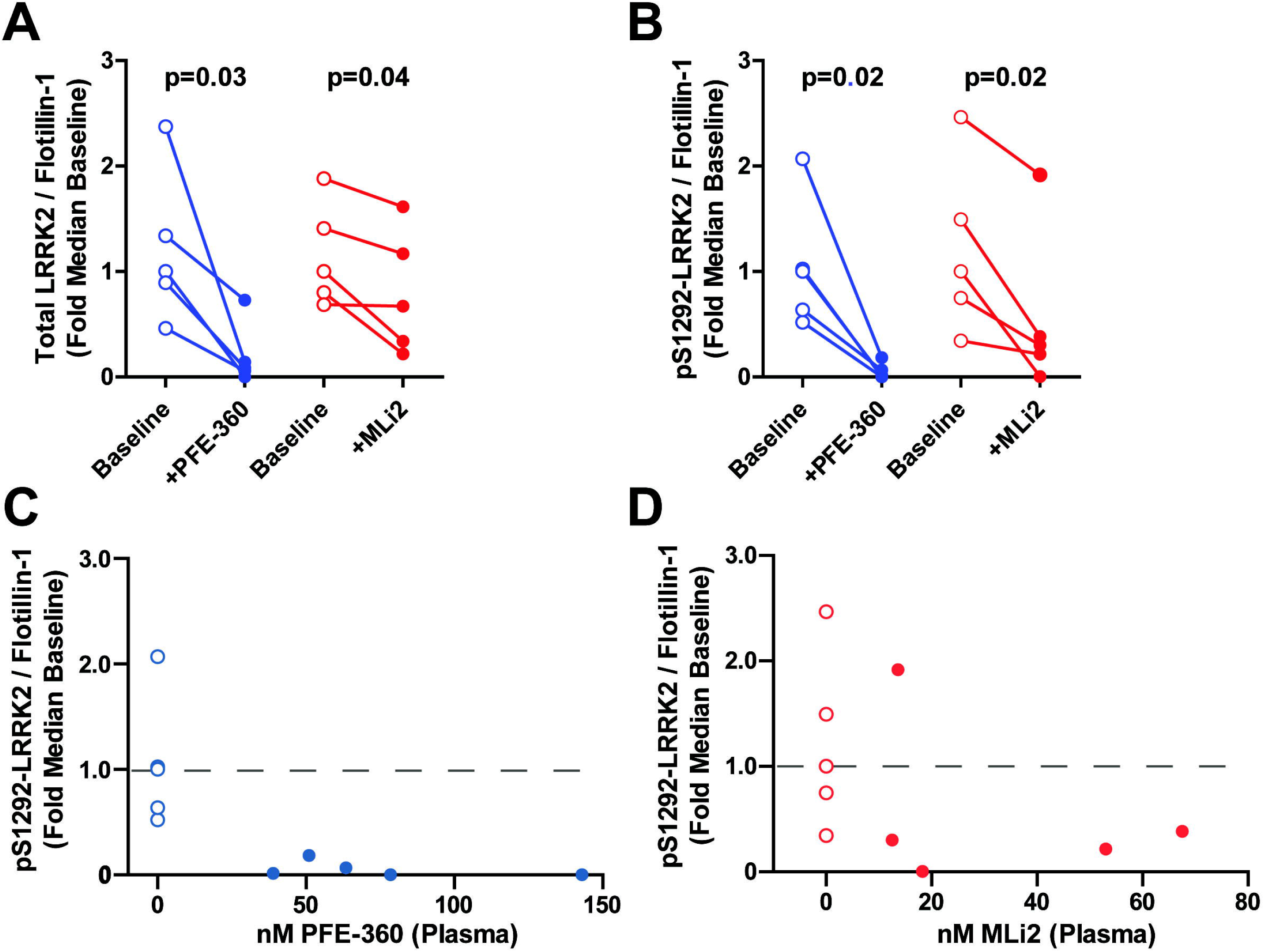
LRRK2 inhibition markers in CSF. Inhibition values for measures of exosomal LRRK2 after five days of PFE-360 (blue) and MLi2 (red) treatment (5 mg kg^-1^ BID). Proteins isolated from CSF exosomes were normalized to the exosome marker Flotillin-1. Fold median baseline values before and post-treatment for **A**. Total LRRK2, **B**. pS1292-LRRK2. Dependent sample t-tests were performed to determine significance between baseline and post-drug exposure measures for PFE-360 and MLi2, respectively. Scatterplots between unbound plasma concentrations of **C**. PFE-360 (nM) or **D**. MLi2 (nM) and inhibition of pS1292-LRRK2/Total LRRK2. The dashed grey lines represent the median value with no drug. Open circles were used to represent baseline levels of each NHP. *p*-values were calculated using a paired t-test.

Previous studies demonstrated that the phospholipid di-22:6-BMP, usually strongly elevated in phospholipidosis (Tengstrand et al., 2010;Liu et al., 2014;Tengstrand et al., 2019), is reduced in the urine of LRRK2 knockout mice and non-human primates treated with the LRRK2 inhibitor GNE-7915 (Fuji et al., 2015). We wondered whether the same effect could be obtained in exosome-depleted urine used for simultaneous measures of exosomal-LRRK2 protein. We measured di-22:6-BMP levels normalized to urine creatinine in the exosome-depleted urine, with baseline concentrations of di-22:6-BMP equal to 12.8 ± 9.18 ng per mg of creatinine. One macaque demonstrated very high baseline levels of di-22:6-BMP in the MLi2-treated group (∼100 ng, Figure 2I). With LRRK2 inhibitor treatment, di-22:6-BMP levels in the exosome-depleted urine dropped to near undetectable levels in the PFE-360 condition (Mdn_PFE-360_=0.05, IQR= 0.01- 0.07, *t*(4)= 2.70, *p*=.05). In the MLi2 group, four out of five macaques showed di-22:6-BMP reduction (Figure 2I). The last macaque had very low levels of di-22:6-BMP at baseline (∼1 ng) and thus showed no further depletion with LRRK2 kinase inhibition. This macaque also had the lowest concentration of MLi2 in plasma at the time of collection (12.5 nM) as well as the least reduction in pS935-LRRK2 in PBMCs in the group (Figure 1B). Exclusion of this macaque from the group revealed a similar trend of reduced di-22:6-BMP in exosome-depleted urine with MLi2 treatment (Figure 2I). Due to the limited amount of urine that could be collected for exosome isolation (i.e., compared to clinical trials), concentrations of di-22:6-BMP were not measured in the total urine specimens (i.e., prior to exosome depletion), in the isolated urine exosomes, or adjusted by exosome-depletion efficiency in individual samples. Overall, reduced di-22:6-BMP was associated with LRRK2 inhibitor treatment.

### Cerebrospinal fluid markers of LRRK2 kinase inhibition

Our previous work in biobanked human CSF demonstrated that LRRK2 protein and pS1292-LRRK2 protein can be readily measured in exosome fractions, but not Rab10 protein, in our assays (Wang et al., 2017). In exosomes isolated from human CSF, LRRK2 autophosphorylation at the serine 1292 residue is much higher (∼5-fold increased on average) as compared to urinary exosomes taken at near the same time from the same subjects (Wang et al., 2017). In macaque CSF exosomes, total LRRK2 levels diminished with PFE-360 treatment (Mdn_PFE-360_=0.09, IQR= 0.03- 0.44, *t*(4)= 3.14, *p*=.03), as well as pS1292-LRRK2 levels (Mdn_PFE-360_=0.01, IQR= 0.001- 0.13, *t*(4)= 4.07, *p*=.02). Both markers were normalized to the exosome housekeeping protein Flotillin-1 (Figure 3A, B and Supplemental Figure 4A).

**Figure 4.**
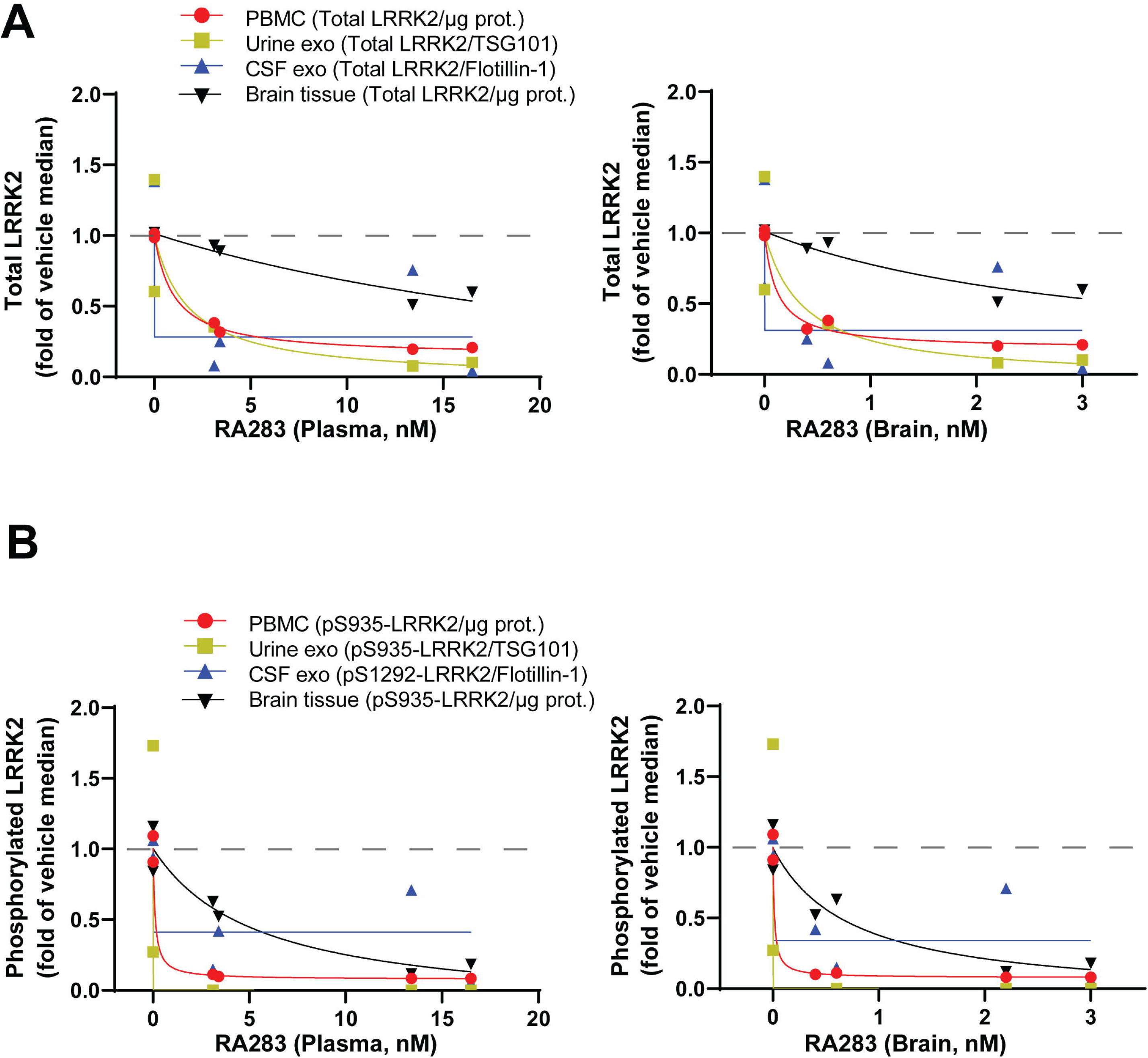
LRRK2 inhibition in RA283 treated NHPs. Inhibition values for measures of LRRK2 in PBMCs, urinary exosomes, CSF exosomes, and brain tissue (frontal cortex) after seven days of low dose (10 mg kg^-1^, 3.4 nM and 3.1 nM) and high dose (30 mg kg^-1^, 13.4 nM and 16.5 nM) RA283 treatment. Biofluids and brain tissue were procured from low dose and high dose NHPs two and six hours after the final dose, respectively. Samples were also collected from vehicle-control NHPs (0 nM RA283). Scatterplots between unbound plasma concentrations of RA283 and **A**. Total LRRK2 and **B**. phosphorylated LRRK2 in PBMCs, urinary exosomes, CSF exosomes, and brain tissue (frontal cortex). One missing sample in urine from a low-dose treated monkey. The dashed grey lines represent the median value with no drug.

Following MLi2 treatment, total LRK2 levels in CSF exosomes were reduced by 35% compared to baseline (Mdn_MLi2_=0.67, IQR= 0.28- 1.39, *t*(4)= 2.98, *p*=.04, Figure 3A). Treatment with MLi2 revealed an exposure-sensitive decrease in pS1292-LRRK2 (Mdn_MLi2_=0.30, IQR= 0.11- 1.51, *t*(4)= 3.56, *p*=.02, Figure 3B, D and Supplemental Figure 4C). Because of the decrease observed with both inhibitors in total LRRK2 protein, ratios of pS1292-LRRK2 to total LRRK2 protein were not calculated. In CSF exosome lysates from all samples, pS935-LRRK2, pT73-Rab10, and total Rab10 could not reliably be detected or measured, presumably due to lower-abundance as compared to urine exosome and PBMC samples. The levels of di-22:6-BMP in the exosome-depleted CSF were not changed with PFE-360 treatment or with MLi2 treatment (Supplemental Figure 4B, D). However, the reliability of the results could be confounded due to variable exosome depletion in individual samples. As with urine analyses, concentrations of di-22:6-BMP were not measured in the total CSF specimens (i.e., prior to exosome depletion) or in the isolated CSF exosomes due to CSF volume limitations in this study.

### LRRK2 kinase inhibition lowers exosome LRRK2 in CSF

In past work with LRRK2 kinase inhibitors in rats and mice, different levels of LRRK2 inhibition were observed with different inhibitors that otherwise possessed very similar drug properties. For example, PFE-360 treatment reduced total LRRK2 levels in brain tissue at much lower concentrations than MLi2, suggesting a drug-dependent target engagement efficacy (Kelly et al., 2018). To better resolve the relationship between brain LRRK2 kinase inhibition and LRRK2 secretion in urinary and CSF exosomes, we tested a third structurally distinct LRRK2 kinase inhibitor, RA283, in a low-dose group and a high-dose group (10 and 30 mg kg^-1^, respectively) compared to a vehicle only-treated group. Macaques were treated (vehicle, low-dose, or high-dose) for a week and sacrificed for the analysis of brain tissue collected at the same time as blood, urine, and CSF to explore the relationship of these fluid markers with inhibition of LRRK2 in brain tissue.

Similar to PFE-360 treatments, RA283 treatment reduced total exosome LRRK2 protein levels in the CSF in three of four treated macaques (Figure 4 and Supplemental Figure 5). Analysis of PBMCs showed a strong reduction of total LRRK2 levels in both low-dose and high-dose treated animals compared to vehicle-control animals (Figure 4A and Supplemental Figure 6). pS935-LRRK2 levels were nearly ablated at unbound drug plasma concentrations that ranged from ∼3 nM to 17 nM (Figure 4B). Consistent with PFE-360 and MLi2 results, a near complete ablation of total LRRK2 protein levels was noted in urinary exosomes at higher plasma drug concentrations (Figure 4A). Evaluation of cortical brain tissue revealed lower total-LRRK2 protein levels (compared to vehicle treated) only in the high-dose animals (Figure 4A). It is worth noting that both doses led to reduced pS935-LRRK2 levels in brain tissue (∼43% at lowest dose and ∼87% at highest dose, Figure 4A). Thus, these results demonstrate that even partial inhibition of LRRK2 kinase activity in brain tissue can lead to a near-complete ablation of total LRRK2 protein levels (and pS1292-LRRK2 protein) in CSF exosomes. These results are consistent with past studies in cell culture suggesting LRRK2-protein release in exosomes is sensitive to kinase inhibition potentially through blocking chaperone activity of 14-3-3 (Fraser et al., 2013;Wang et al., 2019). Such reduction of total LRRK2 protein in the exosome fraction can occur before the total LRRK2 protein in the brain is completely diminished.

**Figure 5.**
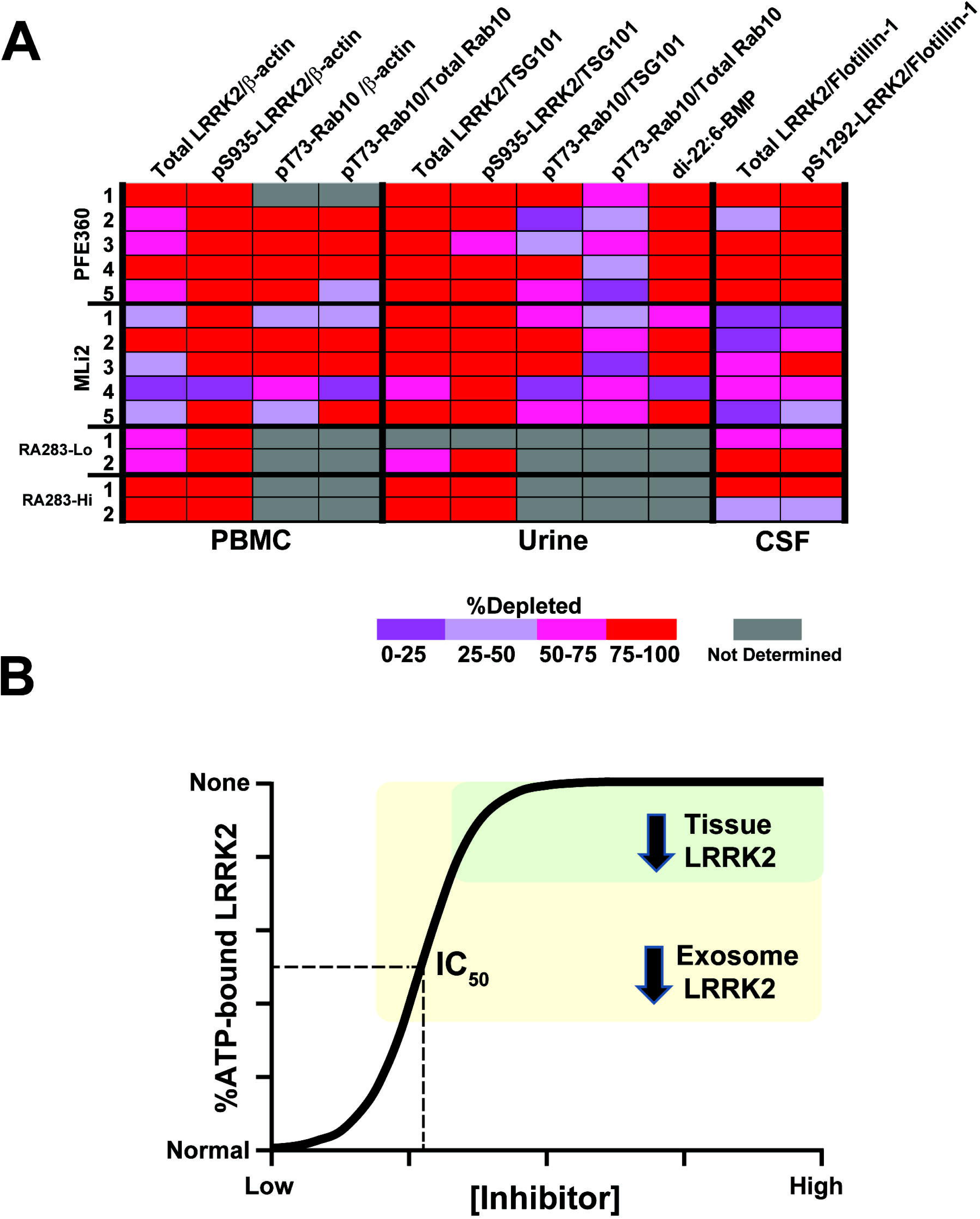
Summary of LRRK2 inhibition. **A**. Heat map detailing the percentage depletion (relative to baseline or vehicle controls) for selected orthogonal pharmacodynamic measures of LRRK2 kinase inhibition across biofluids. Each row represents observations from a separate animal, with biofluids obtained near the same timepoint. **B**. Diagram illustrating a hypothetical relationship of LRRK2 protein in tissue or biofluids (i.e., in exosomes), with %ATP-bound LRRK2 with increasing concentrations of ATP-competitive small molecule LRRK2 kinase inhibitors. In the model, exosome depletion of LRRK2 occurs initially without lowering tissue levels of total LRRK2 is shown in yellow. The green area highlights higher concentrations of drug and corresponding loss of any ATP-bound LRRK2, resulting in the depletion of LRRK2 protein in both exosomes and tissues.

## Discussion

This study identifies efficacious target-engagement measures for LRRK2 kinase inhibition in biofluids (urine and CSF) of non-human primates using three distinct small-molecules (PFE-360, MLi2, and RA283). Our results highlight the variabilities encountered in the different pharmacodynamic measures for LRRK2 inhibition at both baseline (or vehicle-treatment) as well as in response to different plasma levels of drug. All three molecules have very high potency in blocking LRRK2 kinase activity in NHPs and excellent drug-properties for the study. Figure 5A highlights, in individual animals in the study, the effect of the drugs on baseline or vehicle measures of the different pharmacodynamic markers in the study. Differences between the target engagement profiles emerged between the responses in PBMCs, urine, and CSF. While the current standard in the field may be to determine the ratio of phospho-LRRK2 protein to total LRRK2 protein to estimate LRRK2 kinase inhibition, our results here demonstrate that this measure may be critically confounded due to variable decreases in total LRRK2 that can be differentially influenced by distinct classes of small molecules. Instead, we identified housekeeping proteins that may be effective alternatives to ratios of phospho-to total protein. A trend emerged here, consistent with our past studies in transfected cell lines (Fraser et al., 2013), and mice and rats (Kelly et al., 2018), where exosome levels of total LRRK2 protein diminish prior to LRRK2-inhibitor induced decreases in brain tissue (graphically summarized in Figure 5B). In practice, determining LRRK2 inhibitor target engagement in CSF-derived exosomes revealed a similar profile of total LRRK2 protein reduction to that of the autophosphorylated LRRK2 protein (results summarized in Figure 5A).

In past studies, we, and others, demonstrated elevated LRRK2-autophosphorylation and LRRK2 protein levels associated with PD susceptibility and LRRK2 pathogenic mutations (Fraser et al., 2016;Wang et al., 2017). With high endogenous LRRK2 expression in some types of innate immune cells, PBMCs have been championed as a source for pharmacodynamic biomarkers related to LRRK2 inhibition (Thirstrup et al., 2017), and are currently utilized in clinical trials. However, measures in PBMCs alone may poorly predict LRRK2 inhibition in the brain and other tissues, for example with drugs that are poorly brain penetrant or are rapidly eliminated. PBMCs also rapidly turn over and transiently respond to diverse stimuli unrelated to the inhibitor treatment, affecting LRRK2 expression and phosphorylation. Further, our previous study suggests that the efficiency of LRRK2 inhibition is tissue specific and brain LRRK2 protein may be more resilient to inhibition compared to other tissues (Kelly et al., 2018). Ideally, we suggest that a panel of predictive markers that assess LRRK2 inhibition and pharmacodynamic responses across the body will better highlight the most efficacious drugs and dosage regimens. Past studies with drugs like azidothymidine demonstrate that measurements of drug concentration in CSF do not predict brain penetration of the drug and target engagement in the brain (Dykstra et al., 1993;Pardridge, 2011). We suggest that measuring CSF levels of LRRK2 protein in exosomes could contribute to establishing brain penetration in early phase clinical trials (e.g., acute safety studies).

In past work with different LRRK2 pharmacodynamic measures, despite robust measures in analyzing other tissues (i.e., brain, lung, kidney (Kelly et al., 2018)), we have been unable to reliably detect LRRK2 or Rab proteins in urine or CSF procured from rats or mice. The limited biofluids volume obtained from rats and mice compared to NHP appear not to be the reason since we would pool biofluids in excess of volumes studied here. While the reasons behind our limited success in rodents are not clear, we were able to establish robust measures in NHP urine and CSF to profile LRRK2 kinase inhibitors. Pharmacodynamic markers in each biofluid included total LRRK2, pS935-LRRK2, pT73-Rab10, and the ratio of pT73-Rab10 to total Rab1 in PBMCs, total LRRK2, pS935-LRRK2, pT73-Rab10, and di-22:6-BMP levels measured in urine, and total LRRK2 and pS1292-LRRK2 measured in CSF exosomes (Figure 5A). Although proteomic studies using mass spectrometry have suggested the presence of Rab proteins in the CSF, we have not yet detected pT73-Rab10 in the exosome fraction using our assay (Chiasserini et al., 2014). It is unclear whether this challenge is due to the low assay sensitivity or limited sample volumes. It is plausible that the inhibition profile of each drug would further change under different dosages or time intervals post-oral gavage, or in a chronic inhibition study such as an efficacy trial in the clinic. Further, this study did not evaluate the possible effects of age or sex on these markers. Nevertheless, we would predict this panel of markers would be capable of identifying drugs that poorly inhibit LRRK2 in the brain but effectively inhibit LRRK2 in the periphery, for example a drug that may cause differential changes in PBMC and urine markers but not in CSF markers. A further critical test of this hypothesis could involve a LRRK2 kinase inhibitor series that can be modified to either efficiently cross the blood brain barrier or be excluded from the brain. It will be critical to the success of new compounds to utilize these types of assays in early phase clinical trials to ensure sufficient target engagement in the brain. Many new compounds have high attrition rates due to presumed lack of target engagement at doses resolved in safety studies (Morgan et al., 2012). As we expect variability to increase in a more heterogenous clinical population, implementation of informative pharmacodynamic panels of markers may help improve drug development outcomes.

Pathogenic mutations in LRRK2 are now recognized as one of the most common genetic causes of neurodegeneration, and these mutations all appear to result in LRRK2 kinase hyper-activation (Cookson, 2010;West, 2015;2017). Emerging studies also suggest a proportion of idiopathic PD that share hyper-activated LRRK2 phenotypes (Gilks et al., 2005;Bliederhaeuser et al., 2016;Fraser et al., 2016;Cook et al., 2017;Di Maio et al., 2018). Several small-molecule LRRK2 kinase inhibitors and anti-sense oligonucleotides have been evaluated for efficacy and safety in various model systems, including non-human primates (Daher et al., 2015;Fell et al., 2015;Fuji et al., 2015;Volpicelli-Daley et al., 2016;Thirstrup et al., 2017;Zhao et al., 2017). Many if not most therapeutics that target central nervous-system indications move to clinical trials without convincing biomarkers for target engagement. Yet, the use of reliable biomarkers has been shown to increase the chance of success in the clinic (Lopes et al., 2015;Wong et al., 2019). As potential LRRK2-targeting therapeutics like small molecule LRRK2 kinase inhibitors and LRRK2-targeting anti-sense oligonucleotides move forward, there is an acute need for validated pharmacodynamic and theragnostic markers to monitor the success of different classes of LRRK2 kinase inhibitors in blocking LRRK2 activity associated with disease. Notably, the protein measurement assays presented here are based on immunoblotting procedures that, while linear and quantitative in our protocols (Wang et al., 2017), do not scale well into clinically available tests. Further assay development will be needed for broad clinical implementation.

Here, our data suggest that measures of pS935-LRRK2 and pT73-Rab10, each normalized to β-actin, represent blood biomarkers of target engagement with lower inter-animal variability than other markers. PBMCs are easily procured in the clinic and provide a reliable resource for measuring LRRK2 inhibition in circulating cells. Several studies have shown high LRRK2 expression in specific immune cell subsets (Gardet et al., 2010;Kubo et al., 2010;Maekawa et al., 2010;Hakimi et al., 2011;Thévenet et al., 2011;West, 2017). Immunophenotyping blood from PD patients shows increased LRRK2 expression in monocytes and B-cells compared to healthy controls (Bliederhaeuser et al., 2016;Cook et al., 2017). *Ex vivo* treatment of PBMCs with LRRK2 kinase inhibitors result in a reduction of the constitutive LRRK2 phosphorylation site pS935-LRRK2 without acute toxicity (Perera et al., 2016). While we can reproduce the reduction (e.g., with MLi2 treatment) or complete ablation (e.g., with PFE-360 treatment) of pS935-LRRK2, total LRRK2 levels are variably affected in PBMCs that may confound the interpretation of the ratio of phosphorylated to unphosphorylated peptide. We observe the same confounding issue with Rab10, presumably due to changes in protein turnover with phosphorylation as well as inducible expression in immune cell subtypes that may change with kinase inhibition over time, at least in NHPs.

Urine can be obtained non-invasively and in higher abundance than other biofluids, at least in humans (Wang et al., 2017). Our recent proteomics studies suggest urinary exosomes harbor cytosolic proteins that demarcate organs across the body without enrichment of kidney proteins (Wang et al., 2019). Exosomal LRRK2 is both dimerized and phosphorylated, mimicking cytosolic active isoforms (Sen et al., 2009;Deng et al., 2011;Fraser et al., 2013). LRRK2 kinase inhibitors reduce LRRK2 protein in urinary exosomes, possibly as a result of both disrupted secretion and decreased LRRK2 level in the peripheral. pT73-Rab10 levels also decrease, despite of its variable total levels. Di-22:6-BMP, a biomarker of lysosomal dysregulation, was reduced in exosome-depleted urine with LRRK2 inhibitor treatment. These results were consistent with past observations of other structurally distinct LRRK2 kinase inhibitors in NHPs (Fuji et al., 2015). The observed decrease in di-22:6-BMP was similar to that observed in LRRK2 KO mice, and shown to decrease with pS935-LRRK2 and pT73-Rab10 in lysates from brain, kidney, lung, and PBMCs (Steger et al., 2016). A consequence of LRRK2 inhibition would be the decreased phosphorylation of its Rab GTPase substrates and thereby their accumulation in membranes (Steger et al., 2017;Liu et al., 2018). We propose that the shift in membrane-bound Rabs may affect the biogenesis, motility, and/or transport of BMP containing vesicles (e.g., exosomes, secretory lysosomes) outside of the cell, resulting in the observed decrease in BMP in the urine (Alcalay et al., 2020). Although its source(s) remain unclear (i.e., BMPs are found in a variety of tissues and cell types, including brain), reported changes in urinary di-22:6-BMP appear to reflect a LRRK2-driven effect on lysosomal and secretory biology (Baptista et al., 2013).

However, in the brain, our data suggest that LRRK2 inhibition manifests in biofluids first with the reduction of LRRK2 protein that is secreted in exosomes and, at higher drug doses or subsequent to a strong sustained exposure, manifests with reductions in total cellular levels of LRRK2 protein (illustrated in Figure 5B), consistent with our previous observations in cell lines (Fraser et al., 2013). Mechanistically, we propose that 14-3-3 chaperones LRRK2 protein to multivesicular bodies that export LRRK2 in exosomes, and this interaction becomes rapidly disrupted at lower doses of LRRK2 kinase inhibitors than those required for destabilizing LRRK2 protein in total. Our data here show that this relationship, LRRK2 secretion and inhibitor-induced decreases of protein, can be harnessed for effective LRRK2 target engagement assays in blood, urine, and CSF. Overall, we hope these results help provide a path to establish on-target effects of LRRK2-targeting investigational therapeutics in both the periphery and the CNS in clinical populations to improve the chances of a successful clinical trial.

## Conclusions

In past studies, pharmacodynamic markers for LRRK2 inhibition have focused on measures of the ratio of pS935-LRRK2 to total LRRK2 protein and the ratio of pT73-Rab10 to total Rab10 protein in protein lysates from different tissues. More recently, reduced levels of di-22:6-bis(monoacylglycerol) phosphate in urine have been observed with LRRK2 kinase inhibition. Focusing on clinically available biofluids in non-human primates (NHPs), we evaluate different combinations of pharmacodynamic markers for LRRK2 inhibition in urine and CSF and how these different markers compare to LRRK2 inhibition measured in blood at the same time. Quantitative assessments for LRRK2 inhibition in urine and CSF, including the inhibition of LRRK2 secretion in exosomes, provide an inclusive evaluation of LRRK2-targeting therapeutics across the body. These results demonstrate the utility of readily procured biofluids (i.e., blood, urine, and CSF) in measuring LRRK2 inhibition in both the brain and periphery. A panel of biomarkers, such as those utilized in this study, may distinguish LRRK2-directed therapeutics that successfully engage LRRK2 in the brain versus those that skew towards peripheral LRRK2 inhibition.

## Author Contributions

All authors participated in study design, data analysis, drafting, and correction of this manuscript.

## Competing Interests

S.W., K.K., and A.B.W. declare no competing conflicts of interest. L.D., S.B., and N.S. are employees of Sanofi and have shares in Sanofi. J.B.K. and J.M.B. are employees of Atuka and have shares in Atuka Inc. F.H. and E.T. are employed by Nexctea which holds patent rights to the di-22:6-BMP biomarker for neurological diseases involving lysosomal dysfunction (US 8,313,949, Japan 5,702,363, and Europe EP2419742).

### Acknowledgements

The authors thank Philippe Dugay, Stéphanie Eyquem and Isabel A. Lefevre (Sanofi, Rare and Neurologic Diseases Research), Manuela Pereira and Madeleine Coimbra (Sanofi, DMPK).

## Data Availability

The datasets generated and/or analyzed during the current study are available from the corresponding author on reasonable request.

## Additional Information

A supplementary figure set is available for this paper.

## Abbreviations

PD: Parkinson’s disease
LRRK2: Leucine-rich repeat kinase 2 NHPs: Non-human primates
di-22:6-BMP: di-docosahexaenoyl (22:6) bis (monoacylglycerol) phosphate CSF: Cerebrospinal fluid
ATP: adenosine-tri phosphate

